# Edge communities in functional brain networks reveal heterogeneous, overlapping organization across the human lifespan

**DOI:** 10.1101/2025.09.29.679277

**Authors:** Youngheun Jo, Evgeny Jenya Chumin, Richard F. Betzel

**Affiliations:** Department of Psychiatry, University of Michigan, Ann Arbor, MI; Indiana University Alzheimer’s Disease Research Center, Indiana University School of Medicine, Indianapolis, IN; Center for Neuroimaging, Indiana University School of Medicine, Indianapolis, IN; Medical Imaging Research Center, Indiana University School of Medicine, Indianapolis, IN; Department of Neuroscience, University of Minnesota, Minneapolis, MN; Masonic Institute for the Developing Brain, University of Minnesota, Minneapolis, MN

## Abstract

Understanding changes in functional brain organization and their implications in development and aging is one of the central questions in neuroscience. In this study, we used an edge-centric approach to examine cross-sectional differences in functional brain network organization across the human lifespan using resting state functional MRI data from the Nathan Kline Institute - Rockland Sample dataset. By creating edge time series – a framewise multiplication of nodal time series – and clustering them based on their temporal similarities, we were able to identify clusters of edges instead of nodes. This method naturally allows multiple community affiliations per node (brain region), providing a nuanced perspective on network participation compared to conventional hard-partition approaches. To do so, we created age-neutral templates of edge communities – or “eFC lures” – that, when applied, yielded consistent edge communities across non-overlapping subsamples of data. The communities of edges revealed a trajectory of desegregation with aging, suggested to be linked to neural dedifferentiation of activity and cognitive decline in older adults. Additionally, age group-specific lures significantly enhanced the detection of edge community organization compared to the age-neutral version. Combined, these results offer new insights into the heterogeneous, event cluster-level shifts in brain functional organization as well as underscore the importance of age-targeted analytical frameworks throughout the human lifespan.

## INTRODUCTION

The human brain is a complex system composed of neural elements – cells, populations, brain areas – whose physical connectivity helps shape brain-wide activity patterns [1–3]. The structure of this network and the correlated activity it induces, helps to support myriad human behaviors and higher-level cognitive functions that undergo dynamic changes throughout the human lifespan [4–6]. Investigations into characterizing the complex, higher-order organization of these systems and their dynamics across the human lifespan have been increasing [7, 8].

Using resting state functional magnetic resonance imaging (rsfMRI) data, estimates of functional connectivity between brain regions can be summarized using measures of statistical dependencies – e.g. correlation, coherence, mutual information [9, 10]. Studies using rsfMRI have consistently demonstrated changes in segregation of functional networks – decreases in within-network connectivity and increases in between-network connectivity – with aging [11, 12], while the inverse is observed in development [13]. This change is also highly related to modularity, a measure of communities’ intra- and inter-network connectivity, which decreases with aging [9, 11, 12, 14, 15]. However, previous studies have primarily examined brain network organization using methods that create non-overlapping functional modules in which nodes are only assigned to one module.

While overlapping community detection is not uncommon in network science [16, 17], standard approaches in brain networks primarily focus on creating hard partitions of the brain [18, 19]. Recently, we developed an alternative framework of creating communities of edges though clustering “edge time series”, a framewise multiplication of the nodal time series which is mathematically equivalent to the correlation coefficient when averaged across time [20, 21]. One can cluster the edge time series into groups of edges based on their temporal similarities, which projected back to nodes, have been found with a pervasive overlapping community structure [22].

In this paper, we employ this edge-centric perspective and investigate their overlapping community structures in functional brain networks in subjects spanning the human lifespan [23]. We hypothesized that edge communities can advance the current understanding of functional brain network organization through multiple community affiliations of brain regions rather than hard partitions, providing a more nuanced picture of functional network participation. Specifically, we introduce and compare two types of community detection “lures” (community-level edge connectivity templates) – one age-neutral and the other fitted to specific age groups – to investigate whether age group-specific lures enhance sensitivity to age-related shifts in brain organization.

First, our results demonstrate a robust and consistent structure in edge communities in non-overlapping one-tenths of the subjects in the dataset. Our findings re-veal a heterogeneous pattern in network desegregation in aging and increased stability driven by two dominant communities, with an opposite trend in the smallest community. Furthermore, our results demonstrate that age-group-specific lures for edge communities are more robustly fitted than the age-neutral variant, underscoring the importance of age-targeted approaches for a deeper and more nuanced investigation of functional brain changes throughout the human lifespan.

## RESULTS

### Edge communities are heterogeneous and their lures are consistent across data subsamples

Previous studies of lifespan development and aging have predominantly focused on hard partitions of the human brain into networks or communities estimated from node-based, correlated fMRI BOLD activity – i.e. so-called “functional connectivity”. However, because edge communities allow brain regions to have multiple community affiliations, we hypothesized that an edge-centric perspective could provide a complementary and possibly more nuanced view on the functional organization of the human brain across the lifespan. We tested these hypotheses using data using the Nathan Kline Institute Rockland Sample dataset [23].

To do so, we constructed edge time series following procedures outlined in earlier studies [20, 24] from one tenth (*N* = 58 or 59) of the total dataset (*N* = 585) after subjects were removed with high motion or insufficient data (further details in Materials and Methods). In short, using one tenths of the dataset, we created templates or “lures” of edge communities (Fig.1*a, b*). We then repeated K-means clustering to find the most representative partition across the repeats using the adjusted Rand Index (Fig1*c, d*). We then identified the optimal *K* and the corresponding community labels using sum of squared errors (Fig1*e, f*). These community labels were used to create “eFC lures” which were the eFC values in each community, applied to the held-out nine tenths of the dataset. The eFC lures eliminate the need for aligning community labels across subjects while also creating a “group-representative” partition of edge communities (for further details see sections Creating eFC lures from subsamples and K-means clustering).

**FIG. 1.**
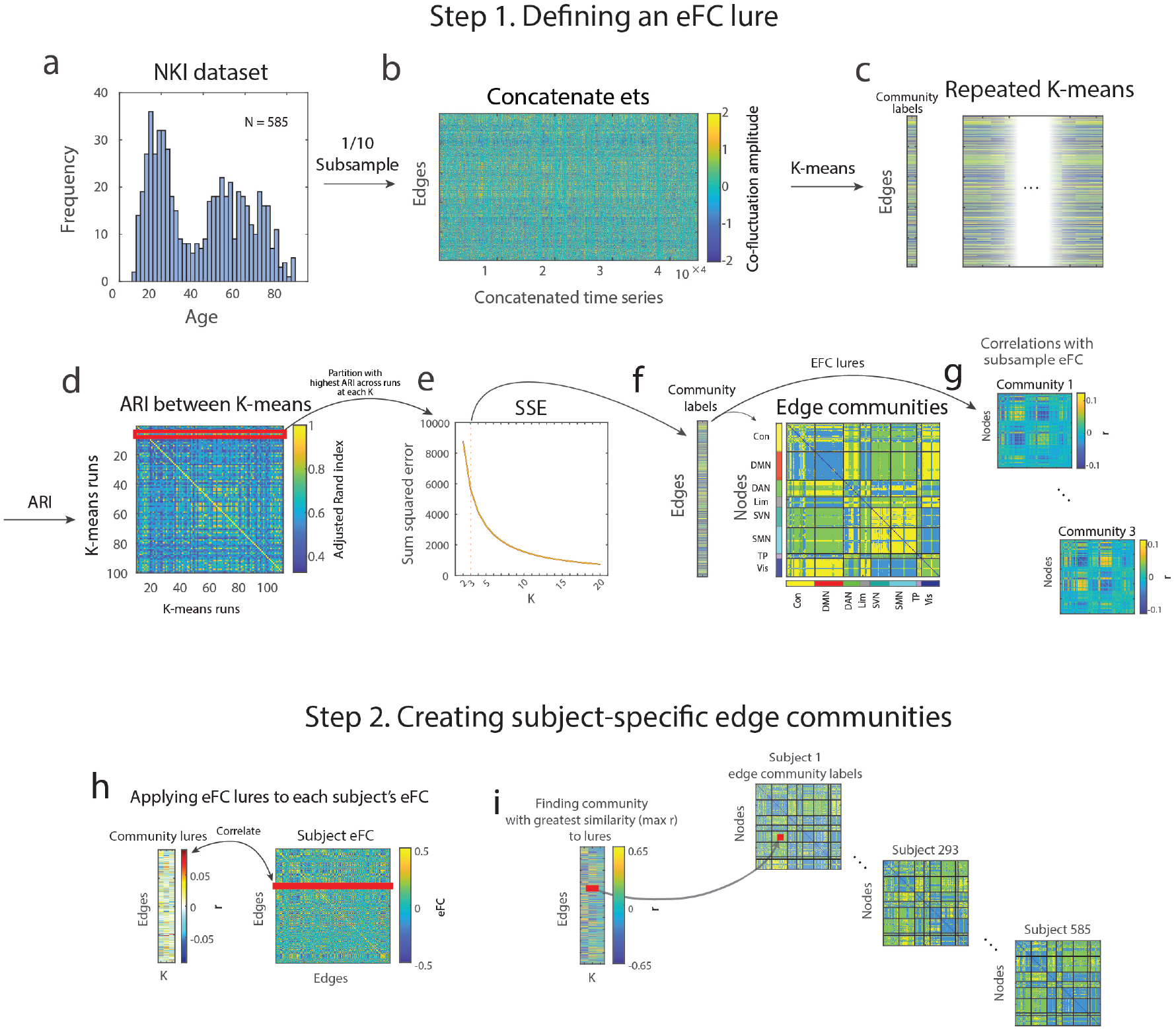
Schematic illustration of creating edge functional connectivity lures for edge communities. (*a*) The age distribution of the NKI dataset. (*b*) Concatenated edge time series of randomly sampled subjects without replacement (1/10th of participants). (*c*) Assigning community labels to each edge using K-means clustering based on edge time series. (*d*) Finding the most representative edge community partition using adjusted rand index (ARI) out of 100 runs of K-means clustering at each *K*. (*e*) Identifying a *K* at which there is a reduction in velocity in change in sum squared error. (*f*) Edge communities at *K* = 3. (*g*) Each community lure’s correlation with eFC. (*h*) Using the community lures, we can apply them to each subject’s eFC and (*i*) find each lure’s best fitted community label for (*j*) each subject. Network abbreviations: Con, control; DMN, default mode; DAN, dorsal attention; Lim, limbic; SVN, salience ventral attention; SMN, somatomotor; TP, temporoparietal; Vis, visual.

Furthermore, we examined the functional organization of the edge communities themselves. As we expected, when examining the functional systems of edge communities and their lures, we found that the community labels and their lures of nodes within-systems were significantly more similar to and less entropic (less random) than nodes spanning canonical functional brain networks (Fig.2*f - i*) [25]. In addition, as hypothesized, we observed the edge communities to be differentially populated with edges spanning functional networks. Community 1 mainly included the edges connecting the control and default mode networks, community 2 with edges spanning the default mode and somatomotor networks, and community 3 with edges connecting the default more and visual networks (Fig.2*j*).

**FIG. 2.**
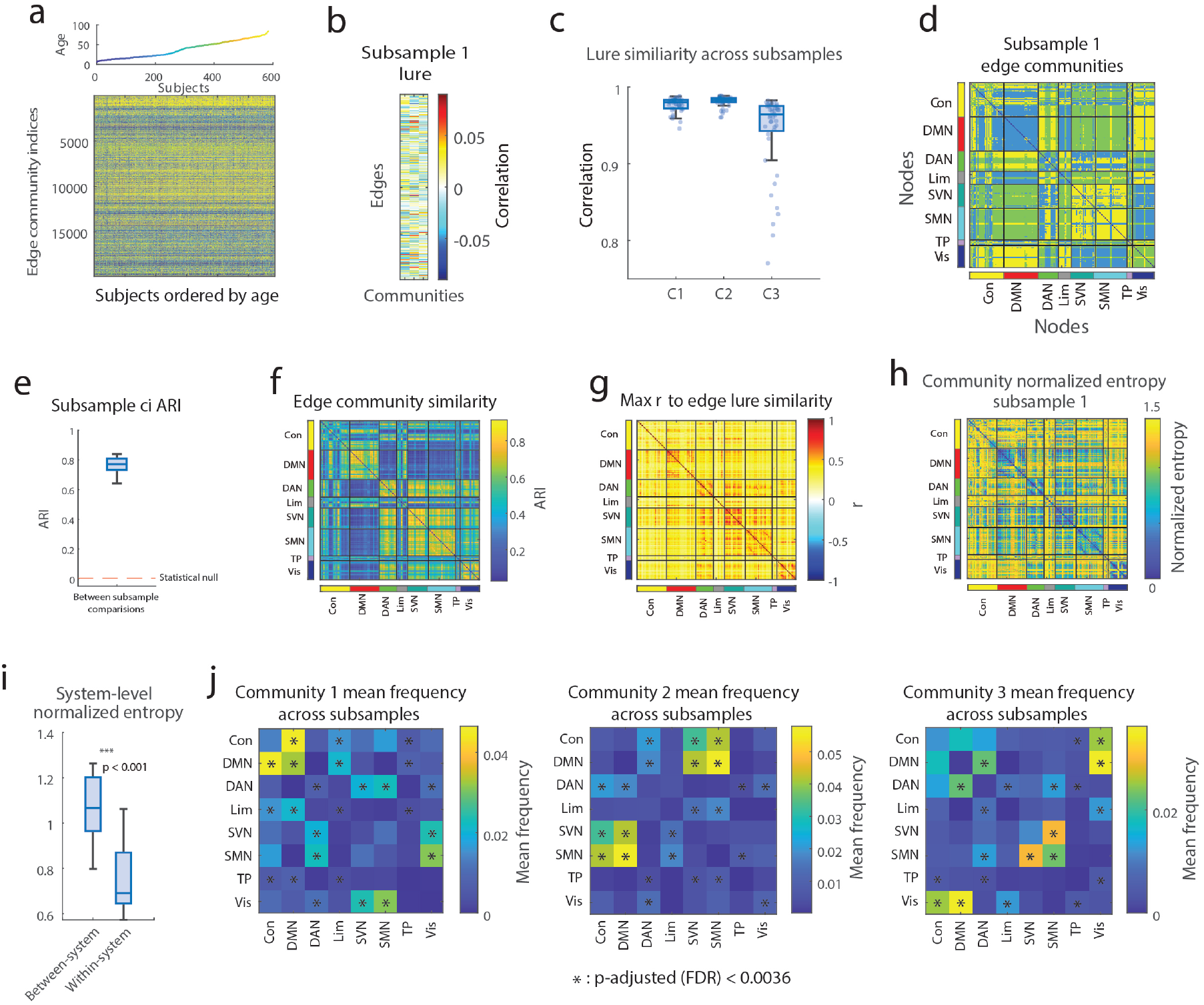
Edge community characteristics. (*a*) Edge community labels across subjects spanning the human lifespan (ages 6 - 84). (*b*) EFC lure created from 1/10th subsample of the dataset. (*c*) Lure similarity across all ten eFC lures created from subsamples comprising 1/10th of the data. (*d*) Edge communities of subsample 1. (*e*) Adjusted Rand index (ARI) for edge communities between subsamples compared to a spin test null. (*f*) Edge community profile similarities between nodes and (*g*) the edge lure similarity profiles between nodes. (*h*) Normalized entropy of edge communities in subsample 1 and (*i*) within-, between-system edges. (*j*) Mean frequency of edges in each community across subsamples (asterisk: significantly more frequent than spin-test null; *padjusted <* 0.0036). Network abbreviations: Con, control; DMN, default mode; DAN, dorsal attention; Lim, limbic; SVN, salience ventral attention; SMN, somatomotor; TP, temporoparietal; Vis, visual.

After creating an edge community lures from one tenths subsample of the dataset, we applied the lures to the remaining 90% of subjects. That is, for each edge, we assigned it the community label of the lure to which its edge connectivity was maximally similar. We were able to identify the edge community labels across held out subjects spanning the human lifespan (Fig.2*a*) from applying subsample 1’s lure (Fig.2*b*). When examining the similarity of the lures themselves, lures were similar across subsamples for each community (Fig.2*c*). Also, the community labels from applying lures were significantly more consistent across subsamples than the spin test null counterpart of 5000 iterations (Fig.2*e*; *p <* 10^−15^).

### Edge community profiles differ across age groups

In the previous section, we found communities of edges to be consistent across non-overlapping, age-neutral subsets of the NKI dataset. Here, we tested whether edge communities, when detected in each subject using age-neutral edge community lures, reveal systematic global differences across the human lifespan.

To do so, we first investigated whether edge communities change frequency with age (Fig.3*a*). The frequencies of communities significantly varied with age, with the larger communities 1 and 2 increasing in frequency with age (*r* = 0.31, *p <* 10^−13^; *r* = 0.10; *p* = 0.015) and the smallest community 3 decreasing with age (*r* = −0.25, *p <* 10^−9^; for other subsamples refer to Fig.S4). We then examined whether edge communities heterogeneously vary across brain regions using a measure called normalized nodal entropy. We found that normalized nodal entropy is significantly lower – edge communities are more variable – in older subjects than in younger subjects (Fig.3*b, c*). Decreases in global normalized nodal entropy were observed across multiple hierarchies of edge community organization (Fig.3*d*; for other *K*s refer to Fig.S6).

**FIG. 3.**
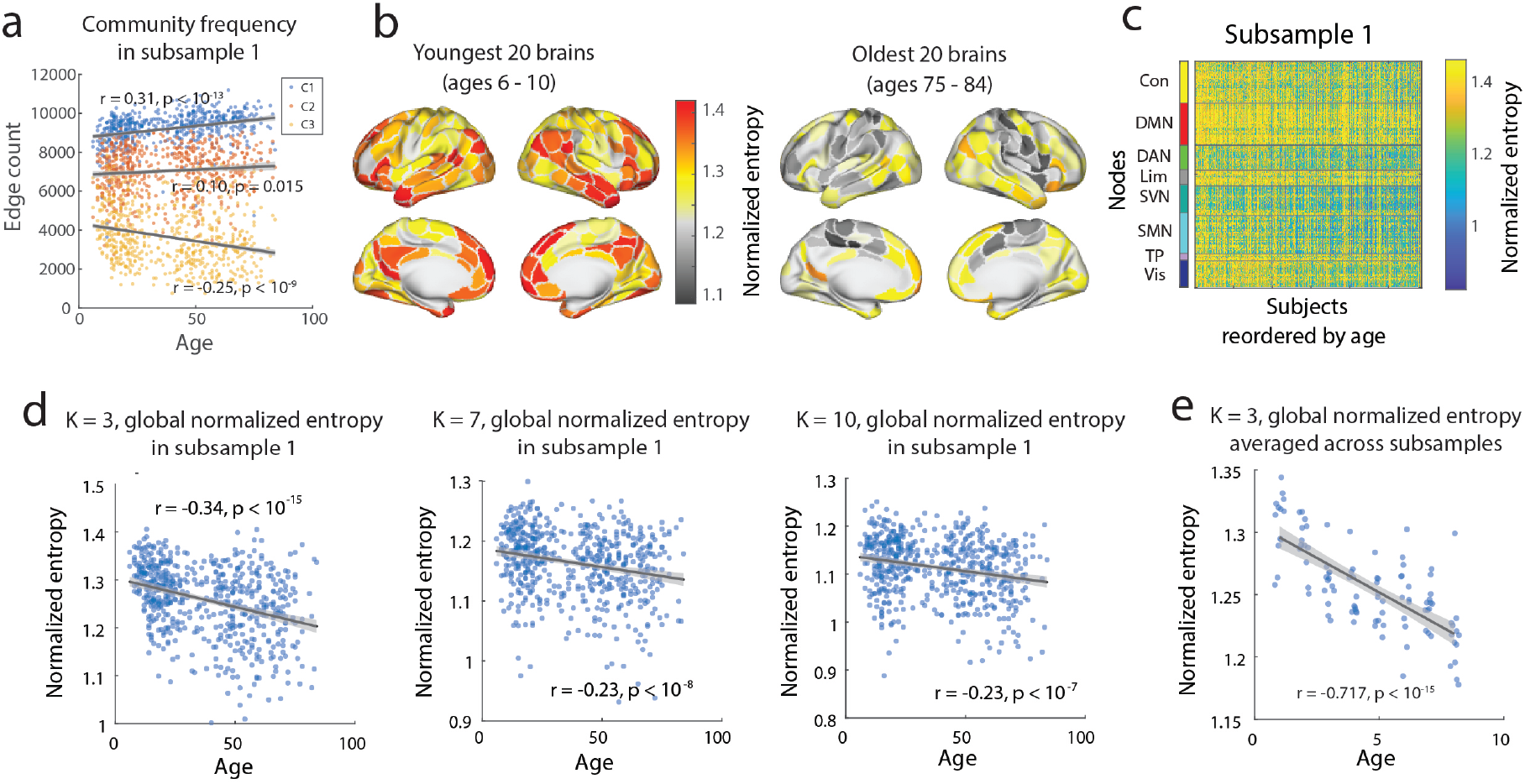
Edge community profiles across the human lifespan. (*a*) Frequency of each community across age in subsample (*b*) Normalized nodal entropy of 20 youngest (left) and 20 oldest subjects’ brains (right) in subsample 1. (*c*) Normalized nodal entropy of each node in subsample 1, subjects ordered by age. (*d*) Global normalized entropy in subsample 1 at K = 3, 7, and 10. (*e*) Global normalized entropy averaged across 10 subsamples of matching numbers of subjects across age groups (age bin = 10 years). Network abbreviations: Con, control; DMN, default mode; DAN, dorsal attention; Lim, limbic; SVN, salience ventral attention; SMN, somatomotor; TP, temporoparietal; Vis, visual.

In addition, due to the differences in the number of subjects in each age bin as in Fig.1*a*, we also created lures of matching numbers of subjects in each age bin (Fig.3*e*). The results from matching numbers of subjects in each age group (age bin = 10 years), averaged across 10 subsamples also revealed a consistent decrease in global normalized entropy with age (*r* = −0.717, *p <* 10^−15^).

### System-level differences in edge communities with age

Previously, we demonstrated that edge communities systematically and globally vary across the human lifespan – with community specific changes in frequency and global normalized entropy to decrease with age. Here, we further tested whether there are system-level differences in edge community organization with age using eFC lures and the best correlation to these lures.

First, we investigated whether there were nodal and system-level differences in correlation to eFC lures – the edge community with the greatest maximum correlation or “max *r*” for each edge – with age. Here, we wanted to choose an appropriate max *r* threshold and using the first derivative of the maximum *r* thresholds, we found that *r* = 0.15 was the optimal across the average of ten subsamples (Fig.4*a*) – which we later used as a threshold for further investigation. Consistent across all thresholds, the counts of all three communities decreased with thresholding without any crossover (Fig.4*b*) with community 1 being the most frequent, and community 3 being the least frequent in the edge count (Fig.4*c*).

**FIG. 4.**
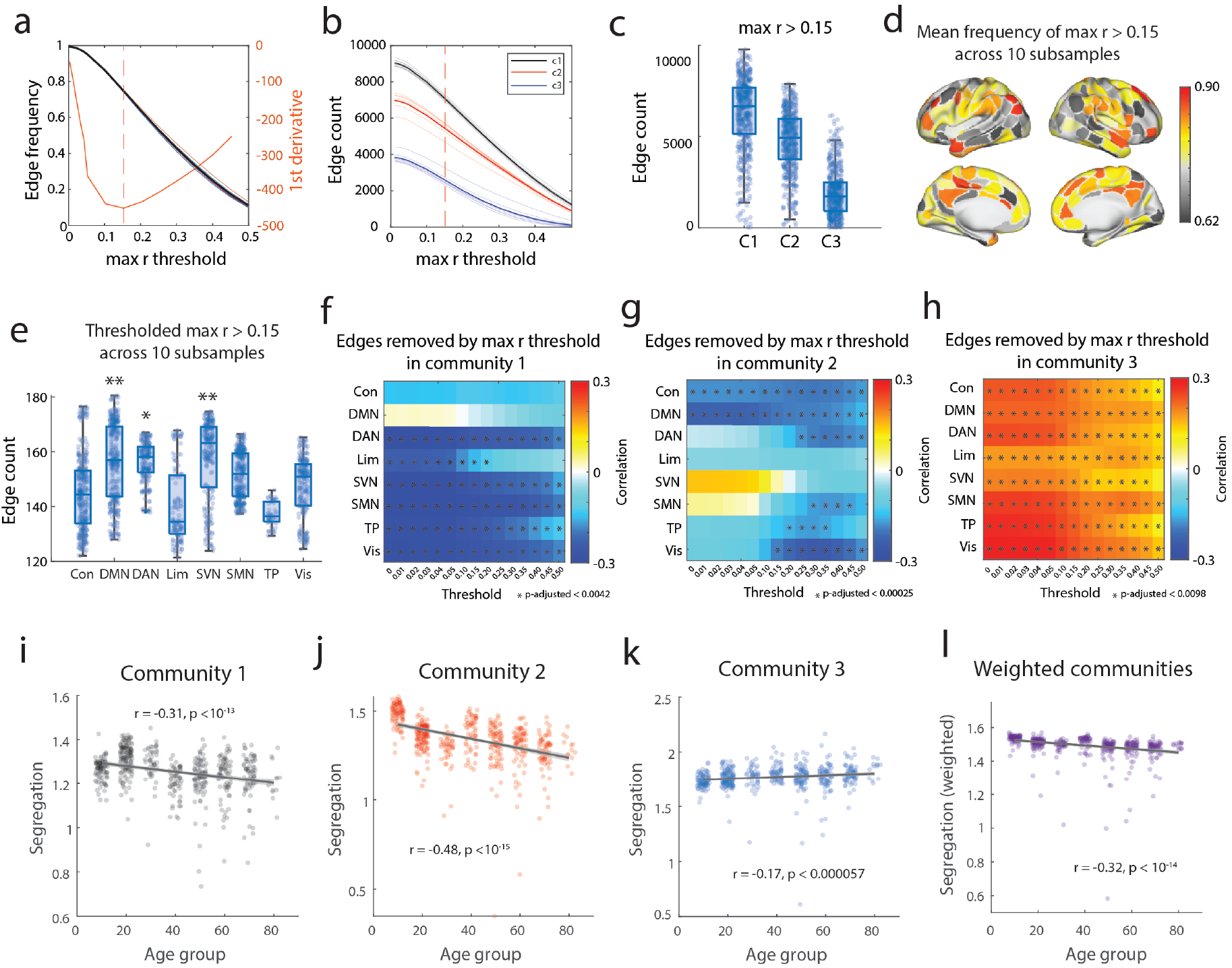
Fit to eFC lures vary with age and system. (*a*) Edge frequencies by thresholding fit to lures by max *r*. (*b*) Edge count by community across different max *r* thresholds. (*c*) Edge count at max *r >* 0.15 for each community. (*d*) Mean frequency of edge counts of max *r >* 0.15 across 10 subsamples. (*e*) System-level frequency of edges at max *r >* 0.15 across 10 subsamples. (*f-h*) Edges removed by max *r* thresholding correlated with age for each community (asterisk: significantly correlated with age above adjusted *p*). (*i-k*) Segregation of each community across age groups and (*l*) segregation weighted by the frequency of each community. Network abbreviations: Con, control; DMN, default mode; DAN, dorsal attention; Lim, limbic; SVN, salience ventral attention; SMN, somatomotor; TP, temporoparietal; Vis, visual.

Next, we tested whether there were system-level heterogeneities in the correlation to the edge community lures. To do so, we zoomed in on the nodal and system-level variance in edges that survived the max threshold of *r* above 0.15 (Fig.4*d, e*). Compared to a spin-test null of 5000 iterations, we found that the edges connecting default mode (DMN), the salience ventral attention (SVN), and the dorsal attention networks (DAN) were significantly more frequent at a maximum threshold of *r >* 0.15 in all 10 subsamples (Fig.4*e*). In addition, when we investigated each community, we found an age dependence in the edges removed through max *r* thresholding. In communities 1 and 2, there were fewer edges removed with age across various max *r* thresholds (*r* = 0 − 0.50; Fig.4*f, g*), whereas more edges were globally removed with age in community 3 (Fig.4*h*). Specifically, in community 1, all networks except the default mode network were significantly less removed in older subjects by thresholding than younger subjects (Fig.4*f*; *p*_*adjusted*_ *<* 0.0042). Edges in community 2 were also significantly less removed in older subjects than in younger subjects in the control, default mode, dorsal attention, somatomotor, temporoparietal, and visual networks (Fig.4*g*; *p*_*adjusted*_ *<* 0.00025). Edges assigned to community 3, the community with the least edges, and the edges removed by thresholding globally positively correlated with age (Fig.4*h*; *p*_*adjusted*_ *<* 0.0098).

Finally, we examined whether edge communities also demonstrated a decrease in segregation with aging, paralleling findings made using nodal functional connectivity [11, 12, 26]. To do so, we tested the segregation of each community, as well as the weighted segregation by each community’s frequencies (Fig.4*i-l*). Community segregation was measured as the within-community edge strengths minus the between-community edge strengths normalized by the within-community edge strengths. As hypothesized, we observed a general desegregation in edge communities with age when weighted by the number of edges in each age group (Fig.4*l, r* = − 0.32, *p <* 10^−14^). However, there were community-level differences – with communities 1 and 2 desegregating with age (*r* = − 0.31, *p <* 10^−13^; *r* = − 0.48, *p <* 10^−15^, respectively) while community 3 became more segregated with age (*r* = −0.32, *p <* 10^−14^).

### Age group-specific lures of edge communities improve lure correlation

Previous studies have reported changes in functional brain organization with age [11, 12, 26]. Most of this work, however, imposes young adult parcellations onto the brains of developmental and aging samples. It is unclear, then, to what extent observed age-related effects might vary if age-specific parcellations were used, instead [25, 27–33]. Here, although we do not vary the parcellation, we examined whether edge communities of rsfMRI data can be better fitted when using age group-specific lures than when using age-neutral lures, and if so, which functional systems exhibit stronger age-group effects compared to than others. The age group-specific lures were created in an identical approach to that of the age-neutral lures, other than using one tenth of the subjects, we used 10 subjects in each age group.

At the edge level, as we hypothesized, subjects were significantly better fitted to age group-specific lures than to age-neutral lures (Fig.5*c*; *p <* 10^−15^) when comparing max *r* values. Furthermore, when applying age group-specific and age-neutral lures, older subjects were found to have a greater nodal max *r* than younger subjects (Fig.5*a, b*). In both age-specific and age-neutral lures, correlation increased with age when applied to each age group (Fig.5*d*; age group-specific lures in orange: *r* = 0.171, *p <* 10^−15^; age-neutral lures in blue: *r* = 0.134, *p <* 10^−15^). Across all age groups, age group-specific lures were better fitted to the corresponding age group than age-neutral lures.

**FIG. 5.**
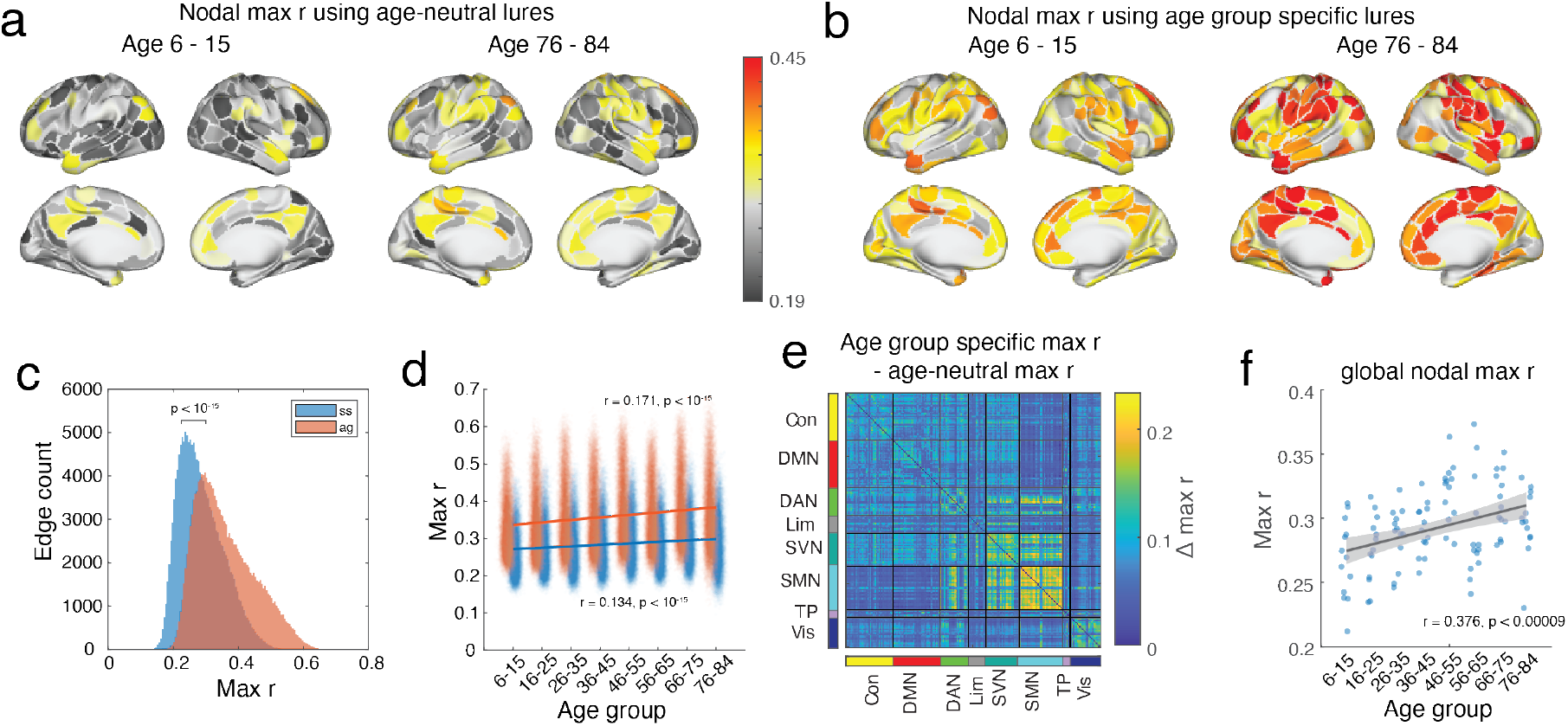
Applying age group specific lures. (*a*) Nodal max *r* of youngest (left) and oldest (right) age groups when applying each age group’s specific lures. (*b*) Nodal max *r* of youngest (left) and oldest (right) age groups when applying each age-neutral lures. (*c*) Max *r* values of edges from age-neutral, subsampled lures (ss; blue) and age group-specific lures (ag; orange; *p <* 10^−15^). (*d*) Max *r* values of edges when applying age group-specific lures to each age group (orange; *r* = 0.171; *p <* 10^−15^) and age-neutral, subsample lures to each age group (blue; *r* = 0.134; *p <* 10^−15^). (*e*) Differences in max *r* between age group-specific minus age-neutral, subsample lures. (*f*) Global nodal max *r* in matching sized age group lures applied to each age group. Network abbreviations: Con, control; DMN, default mode; DAN, dorsal attention; Lim, limbic; SVN, salience ventral attention; SMN, somatomotor; TP, temporoparietal; Vis, visual.

Next, we tested whether the fit to the lures was system-dependent. As we expected, subjects were generally significantly better fitted to age group-specific lures than age-neutral lures (*p*_*adjusted*_ = 0.00002, false discovery rate = 0.01), except for edges connecting nodes within the temporoparietal network (Fig.5*e*), with the greatest differences centering on the somatomotor network. We also tested whether the number of subjects in each age group affects lure correlation with age. When matching the number of subjects in each age group for creating eFC lures and applying the lures to matching numbers of subjects in each age group, we found consistent age-related trends as in Fig.5*d* (Fig.5*f*; *r* = 0.376, *p* = 0.00009).

## DISCUSSION

Seminal findings in the aging literature have demonstrated an age-related trend in neuronal signals to become more diffuse and reduced with age, a process also known as neural dedifferentiation [34]. Similarly, this trend with aging has been demonstrated in recent studies, revealing network desegregation – increased between-network and decreased within-network connectivity – with age [11, 14, 26, 35–39]. However, previous methods have mainly focused on network segregation using “hard partitions” of brain function, which is both biologically and functionally unlikely, limiting a nodes’ role or affiliation to a single function or community [40, 41]. In this study, we used an edge-centric approach that naturally creates overlapping nodal partitions, allowing multiple functional participations of each brain region [20]. Specifically, we created and tested two types of templates or “lures” of edge community detection: one that closely parallels the U.S. demographic distribution and another that is specific for each age group.

We found that, as hypothesized, edge communities are consistent across non-overlapping subsets of the data, with partitions more similar between nodes within than between canonical functional systems [25]. In large, our results also demonstrate an overall age-related desegregation in edge communities, along with heterogeneous, community-specific changes. Specifically, communities 1 and 2 decreased in segregation with age and increased correlation to lures with age, while community 3 increased in segregation with age and showed decreased correlation to lures with age. The changes in global segregation are likely driven by the more frequent, dominant communities 1 and 2 revealing community-dependent dynamics in functional organization with age – which can be eluded when investigating hard nodal partitions alone.

However, our results also varied in part from the existing literature with decreasing nodal and global normalized entropy of edge community partitions across multiple scales of community sizes (*K* = 3, 7, 10). Globally, we reported less variability in community partitions (measured *via* normalized entropy) and increased correlation to lures in older subjects than in younger subjects. In the smallest community 3, however, we reported increases in normalized entropy and decreased correlation to lures with age. These results reveal a heterogeneous and potentially compensatory process between edge communities with dominant communities 1 and 2 increasing in frequency and correlation while decreasing in segregation with age, whereas community 3 decreases in frequency and correlation but increases segregation with age. Combined, the overlapping community structures of the human brain may become canalized, less variable, and less segregated with healthy aging. Our study provides further support for the pervasive overlapping functional organization of the human brain and the overall desegregation with healthy aging.

Although our results highlight age-related patterns of desegregation and segregation within edge communities, caution is warranted when directly comparing these edge-centric measures to conventional, node-based approaches. In an edge-centric framework, a single node may belong to multiple functional communities *via* its edges; thus, “segregation” in the edge domain may reflect fundamentally different organizational properties than node-centric segregation. For instance, an edge community with high internal connectivity (i.e., high segregation from other edge communities) could nonetheless represent between-network or between-community connections at the nodal level. Therefore, interpreting changes in edge community segregation should not be taken as a direct analogue to nodal or canonical network segregation. Instead, these two perspectives - node-centric versus edge-centric — capture different facets of brain organization and should be regarded as complementary approaches rather than being interchangeable. Future work may benefit from systematically comparing these measures and elucidating how differences in functional organization relate to cognition, aging, and clinical outcomes.

Finally, our study further highlights the importance of age group-specific lures over an age-neutral counterpart. Previous studies have drawn caution in applying functional lures derived from healthy young adults to the rest of the healthy population, especially given age-related changes in the brain’s functional architecture [29–33]. Our results demonstrate that when comparing the correlation of an age-neutral lure to that of an age group-specific lure, there are significant differences that vary by the functional system. Globally, age group-specific lures were better fitted to subjects in their respective age groups than the age-neutral lures, except for the temporoparietal system. This may be due to the subject-specific organization or intersubject variabilities of this system rather than differences with age. We also note that while the differences to the two types of lures are global and significant, these differences in correlation are limited. This may in part be due to the age-neutral lures including subjects from all age groups, rather than from lures derived from a small subset of the population (e.g. young healthy adults).

Furthermore, system-level differences in the two types of lures were maximized in the somatomotor areas. Previous studies have demonstrated age-related increases in functional connectivity within motor regions [42, 43]. A greater fit in max *r* for edges within the somato-motor network suggests age-related changes in resting state fMRI data that involve these cortical areas that are better captured *via* age group-specific investigations. These system-level heterogeneities in the correlation to age-neutral versus age group-specific lures can be useful to take into account when applying targeted and narrow (e.g. young healthy adults) functional templates to other age groups.

Recent trends in neuroscience are moving towards more specialized and individualized approaches to enhance the correlation and interpretability of the findings, especially in clinical settings [44–47]. In addition, a dataset and lure of brain function and its organization on a population-general level is necessary to understand inter-population, inter-demographic differences and to maximize representativeness of a population [23]. Previous studies investigating subject-specific edge community structures have reported consistent community profiles within subjects across multiple rsfMRI scans [22] and the entropy of edge communities to be a neural correlate of aging and a moderator of fluid cognition [39]. Future research is required for more nuanced, specialized applications in different age groups and clinical settings to identify heterogeneous shifts and differences in overlapping community structures.

### Limitations and future directions

One of the key limitations of this study is that we used a cross-sectional dataset including subjects spanning the human lifespan. While the NKI dataset includes subjects with demographics representing the US population, the number of subjects across different age groups vary significantly (smallest age group = 13; largest age group = 137); which can impact the number of subjects included in each subsample, as well as the variance in results in each age group. Therefore, as an exploratory analysis, we also matched the number of subjects in each age group when creating and applying lures – in which we also found consistent results with our main findings. Nonetheless, further investigation is required to determine the number of subjects or the scan length required for estimating stable edge community lure in samples that are not “typically aging” or healthy. Another limitation of this study is that our study focused on describing cross-sectional differences in edge community organization, which still leaves questions for mechanisms and drivers in shifts in edge community organization. Future research should investigate the relationship between edge community organization, age, and cognitive functions using longitudinal datasets.

### Conclusion

In this study, we used age-neutral lures of edge communities to examine the functional organization across the human lifespan at the nodal, edge, network, and whole-brain levels. Our results demonstrate a complex and heterogeneous organization of overlapping communities across the human lifespan, which provides a comprehensive examination of the brain’s functional make up and aging process. In addition, our results also highlight the importance of age group-specific lures in investigating the functional organization of human brains – necessitating the development of both age-targeted and population-general lures for improving fMRI-based community estimations.

## MATERIALS AND METHODS

### Dataset

#### Nathan Kline Institute, Rockland Sample

The Nathan Kline Institute Rockland Sample (NKI-RS) dataset consisted of resting state functional magnetic resonance imaging, structural magnetic resonance imaging, as well as diffusion magnetic resonance imaging data from 711 subjects (downloaded December 2016 from the INDI S3 Bucket) of a community sample of participants across the human lifespan. After excluding subjects based on data and metadata completeness and quality control (see Image Quality Control), the final subset used included 585 subjects (63.1% female, age range = 6 - 84). The study was approved by the Nathan Kline Institute Institutional Review Board and Monclair State University Institutional Review Board and informed consent was obtained from all subjects. Subjects were compensated for their participation. A comprehensive description of the imaging parameters can be found online at the NKI website. Briefly, images were collected on a Siemens Magneton Trio with a 12-channel head coil. Subjects underwent three differently parameterized resting state scans, but only one acquisition is used in the present study. The fMRI data was acquired with a gradient-echo planar imaging sequence (TR = 645ms, TE = 30ms, flip angle = 60^°^, 3mm isotropic voxel resolution, multi-band factor = 4). This resting state run lasted approximately 9:41 seconds, with eyes open and instructions to fixate on a cross. Subjects underwent one T1-weighted structural scan (TR = 1900ms, TE = 2.52 ms, flip angle = 9^°^, 1mm isotropic voxel resolution) and one diffusion MRI scan (TR = 2400ms, TE = 85ms, flip angle = 90^°^, 2mm isotropic voxel resolution, 128 diffusion weighted volumes, b-value = 1500s/mm^2^, 9 b = 0 volumes).

### Image Quality Control

The NKI was downloaded in December of 2016 from the INDI S3 Bucket. At the time of download, the dataset consisted of 718 resting state fMRI (“acquisition645”; 634 subjects) 957 T1w (811 subjects), and 914 DWI (771 subjects) images. fMRI images were excluded if greater than 15% of time frames exceeded 0.5mm framewise displacement. Furthermore, fMRI images were excluded if the scan was marked as an outlier (1.5x the inter-quartile range in the adverse direction) in 3 or more of the following quality metric distributions: DVARS standard deviation, DVARS voxel-wise standard deviation, temporal signal-to-noise ratio, framewise displacement mean, AFNI’s outlier ratio, and AFNI’s quality index. Following image quality metric filtering processes excluded 49 fMRI subjects with 585 subjects maintained.

### Image Preprocessing

The fMRI images in the NKI dataset were preprocessed using the fMRIPrep version 1.1.8 [48]. The following description of fMRI preprocessing is based on fMRIPrep’s documentation. This workflow utilizes ANTs (2.1.0), FSL (5.0.9), AFNI (16.2.07), FreeSurfer (6.0.1), nipype [49], and nilearn [50]. T1w images were submitted to FreeSurfer’s cortical reconstruction workflow (version 6.0). The FreeSurfer results were used to skull strip the T1w, which was subsequently aligned to MNI space with 6 degrees of freedom. Functional data was slice time corrected using AFNI’s `3dTshift` and motion corrected using FSL’s `mcflirt`. “Fieldmap-less” distortion was performed by co-registering the functional image to the same-subject T1w with intensity inverted [51] constrained with an average fieldmap template [52], implemented with `antsRegistration`. This was followed by co-registration to the corresponding T1w using boundary-based registration [53] with 9 degrees of freedom, using `bbregister`. Motion correcting transformation, field distortion correcting warp, and BOLD-to-T1w transformation warp were concatenated and applied in a single step using `antsApplyTransforms` using Lanczos interpolation. Frame-wise displacement [54] was calculated for each functional run using Nipype. The first four frames of the BOLD data in the T1w space were discarded. Each T1w was corrected using `N4BiasFieldCorrection` [55] and skull-stripped using `antsBrainExtraction.sh` (using the OASIS template). The ANTs derived brain mask was refined with a custom variation of the method to reconcile ANTs-derived and FreeSurfer-derived segmentations of the cortical gray matter of Mindboggle [56]. Brain tissue segmentation of cerebrospinal fluid (CSF), white matter (WM) and gray matter(GM) was performed on the brain-extracted T1w using `fast` [57].

### Network definition

#### Parcellation

For the NKI resting state fMRI data, the Schaefer 200 parcellation was rendered as a volumentric parcellation in each subject’s anatomical space within the gray matter ribbon. To transfer the parcellation from fsaverage to subject space, FreeSurfer’s `mris_ca_label` function was used in conjunction with a pre-trained Gaussian classifier surface atlas [58] to register cortical surfaces based on individual curvature and sulcal patterns.

#### Functional connectivity

For the fMRI data, each preprocessed BOLD image was linearly detrended, band-pass filtered (0.008-0.08Hz), confound regressed, and standardized using Nilearn’s `signal.clean` function, which removes confounds orthogonally to the temporal filters. The confound regression strategy included six motion estimates, mean signal from the white matter, cerebrospinal fluid, and whole brain mask, derivatives of these previous nine regressors, and squares of these 18 terms. Spike regressors for frames with motion greater than 0.5mm frame-wise displacement were applied. The 36 parameter strategy (with and without spike regression) has been shown to be a relatively effective option to reduce motion-related artifacts [59]. Following these preprocessing operations, the mean signal was acquired for each node in the volumetric anatomical space.

### Edge time series

Following the preprocessing and sampling steps for rsfMRI data described previously, the mean signal was taken at each time frame for each node, forming the nodal time series. The FC between brain regions *i* and *j* is operationalized as a correlation coefficient summarized as the Pearson correlation coefficient as follows:

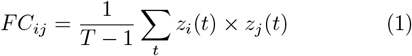

where *z*_*i*_ = [*z*_*i*_(1), …, *z*_*i*_(*T*)] is the vector or z-scored nodal activity from region *i*.

The edge time series for edge *i, j* is calculated by simply omitting the summation and normalization step. In short, edge time series is calculated as follows:

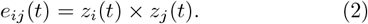

This procedure is repeated for all pairs of nodes resulting in an edge-by-time series matrix. The elements of this matrix encode the moment-by-moment co-fluctuation magnitude of nodes *i* and *j*. A positive value in this 10 co-fluctuation would indicate a simultaneous increase or decrease in the activity of nodes *i* and *j*, whereas a negative value would reflect their opposite direction of activity. Similarly, a magnitude close to zero would indicate that either *i* or *j* had very low levels of activity.

### Creating eFC lures from subsamples

With the edge time series of each non-overlapping sub-sample of the dataset, we cluster edges based on their co-activation patterns between brain regions across time using k-means clustering. Using the optimal number of edge communities, we created lures of edge communities or “eFC lures” which were the eFC values in each community. The held out subjects’ eFCs were then correlated with each edge community lure to find the greatest fitted community with the “max *r*”. This process allows a fast and lure-based edge community detection that also reflects individual-level differences in edge communities.

### K-means clustering

We used a k-means clustering algorithm with Pearson correlation as the distance measure to cluster the edge time series. More specifically, each edge is partitioned as cluster 1 up to cluster *K* based on their edge time series. We acquired the edge partitions for *K* = 2 − 20 with 100 repeats at each *K* and identified the most representative partition at each *K* using the adjusted Rand index. The partition labels in k-means clustering are assigned randomly – therefore, even an identical partition can be labeled differently as follows [*C*1, *C*1, *C*1, *C*2, *C*2, *C*1] or [*C*2, *C*2, *C*2, *C*1, *C*1, *C*2]. Therefore, after acquiring 100 partitions at each *K*, we first identified the most representative partition out of the 100, by calculating the adjusted Rand index between each partition which is calculated as follows with *n*_*ij*_ as the number of elements in the intersection of partition *i* and partition *j* and *a*_*i*_, *b*_*j*_ as the sum of elements in cluster *i* and *j*, respectively:

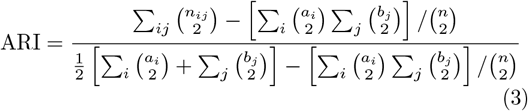

The partition with the highest adjusted Rand index was used to calculate the sum of squared errors across *K* = 2 − 20 which we identified the *K* with the maximum curvature.

After identifying the number of clusters, we realigned the cluster labels of the maximum adjusted Rand index across the ten subsamples so that each subsample will have cluster labels that are maximally similar to the cluster labels in another run. For the alignment, we first identified the most representative partition out of the ten subsamples by calculating the adjusted Rand index between each partition. Next, we realigned the other nine subsamples to the most representative partition using the *matchpairs* function provided in MATLAB that is based on the Hungarian algorithm that aims to minimize total cost - measured as cluster centroid dissimilarity (1 − Correlation coefficient) - of a linear assignment problem. Cluster centroids from each run were then re-aligned to match the centroids of the partition that minimizes the total cost.

### Normalized entropy

Normalized entropy was calculated for the edge community labels as follows where *H*_*P*_ is the entropy of the *i*th node and *p*_*ic*_ is the fraction of node *i*’s connections assigned to edge community *c*:

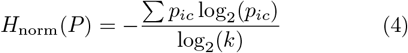

Global normalized entropy was calculated by averaging across all nodal normalized nodal entropies.

## Author Contributions

YJ, EC, and RFB conceived of study and YJ carried out all analyses, and generated figures. EC contributed to the project *via* discussion. All authors helped revise and write the submitted manuscript.

## Data and code Availability

All imaging data come from publicly-available, open-access repositories. Human connectome project data can be accessed *via* https://db.humanconnectome.org/app/template/Login. after signing a data use agreement. Midnight scan club data can be accessed *via* OpenfMRI at https://openfmri.org/dataset/ds000224/.

---

**FIG. S1.**
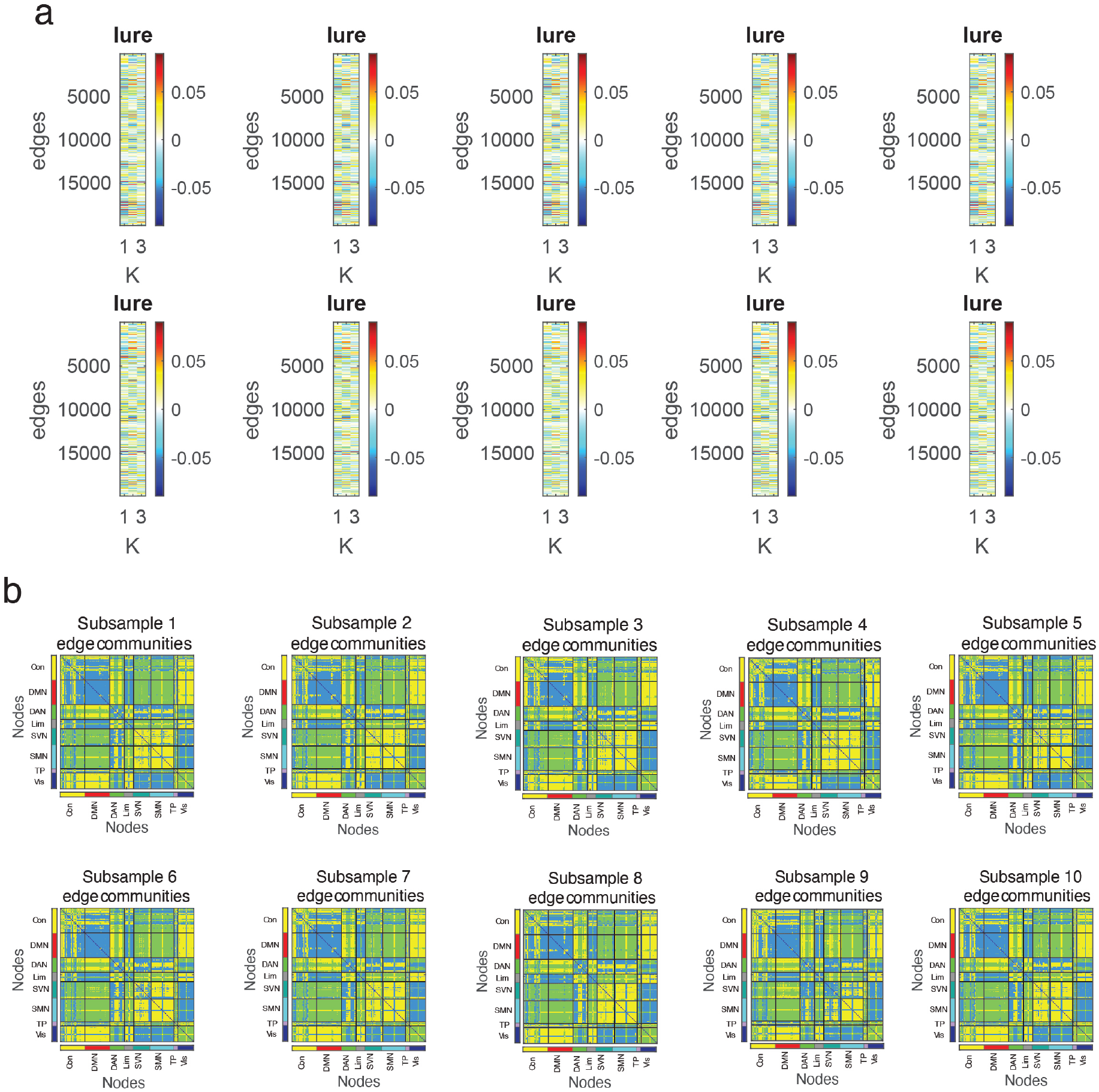
EFC lures and edge communities across ten non-overlapping subsamples of the NKI dataset. (*a*) EFC lures in each subsample and (*b*) the edge communities applied to the held out subjects.

**FIG. S2.**
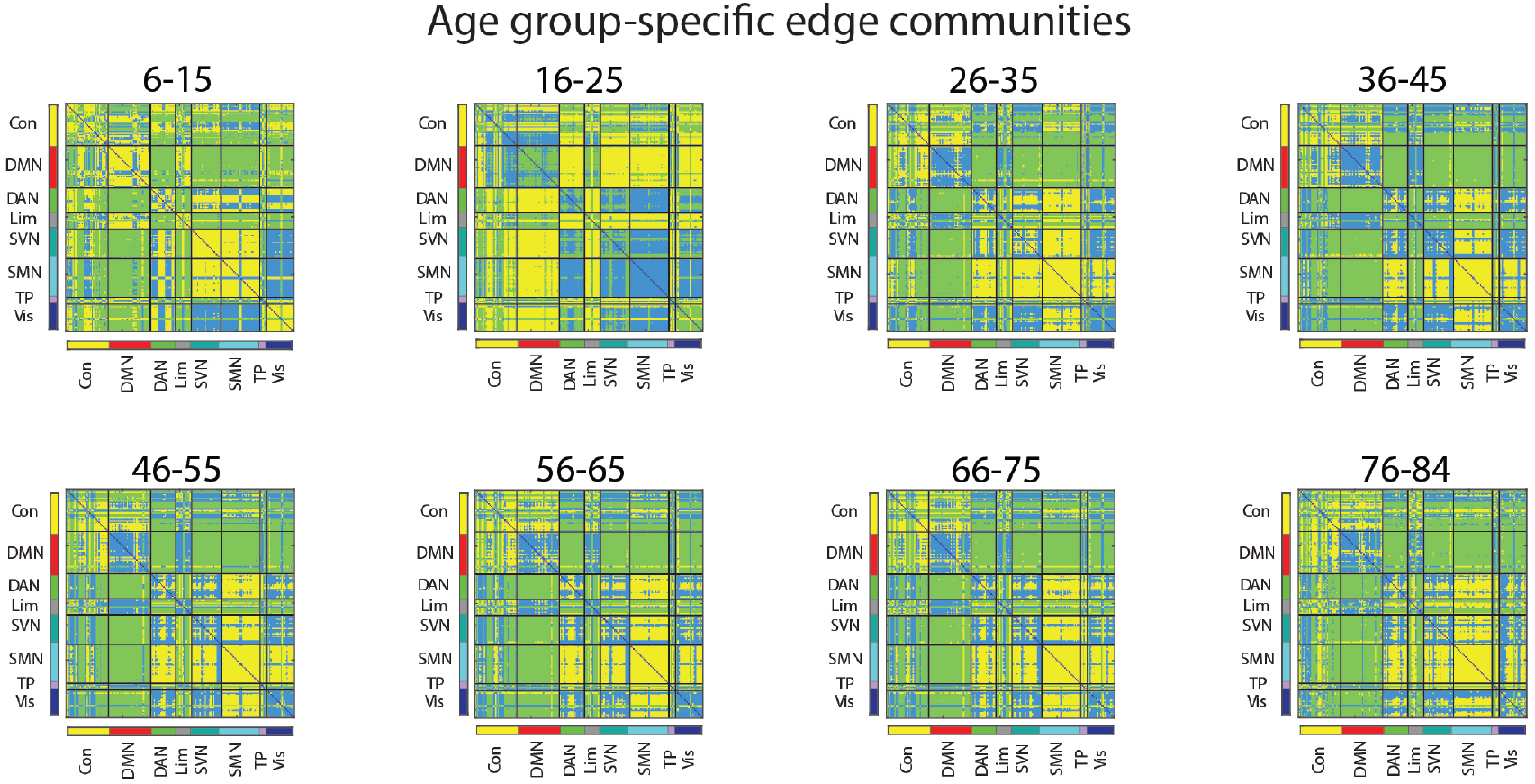
Age group-specific edge communities. Edge communities of each age group (age bin = 10 years) realigned to the partitions of the subsamples using the Hungarian algorithm.

**FIG. S3.**
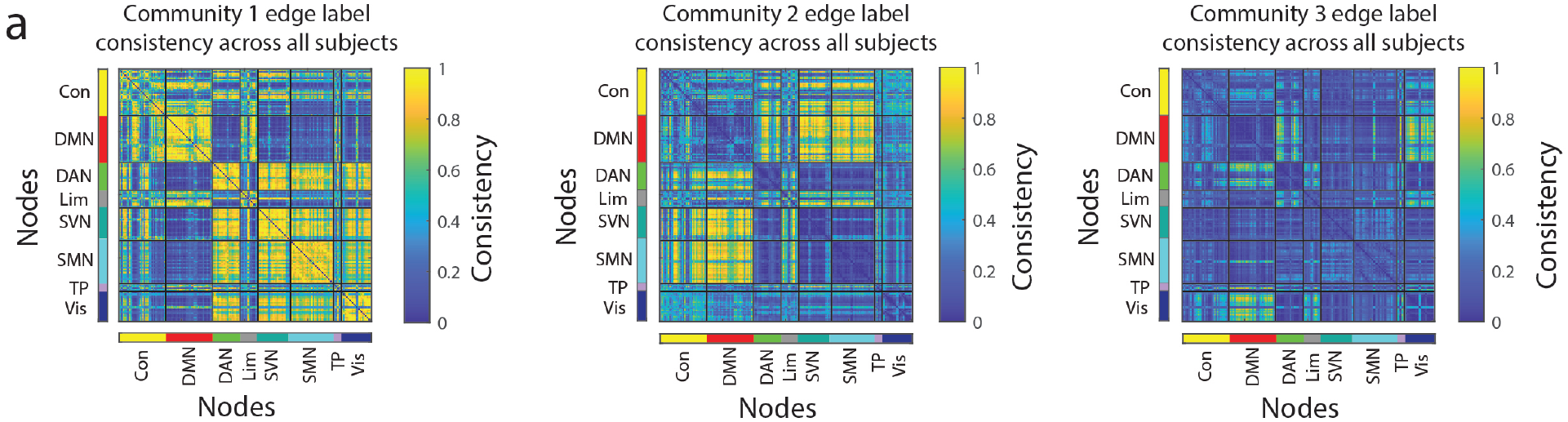
Edge community consistency across all subjects. Community label consistency for each edge across all subjects in each community, averaged across 10 subsamples.

**FIG. S4.**
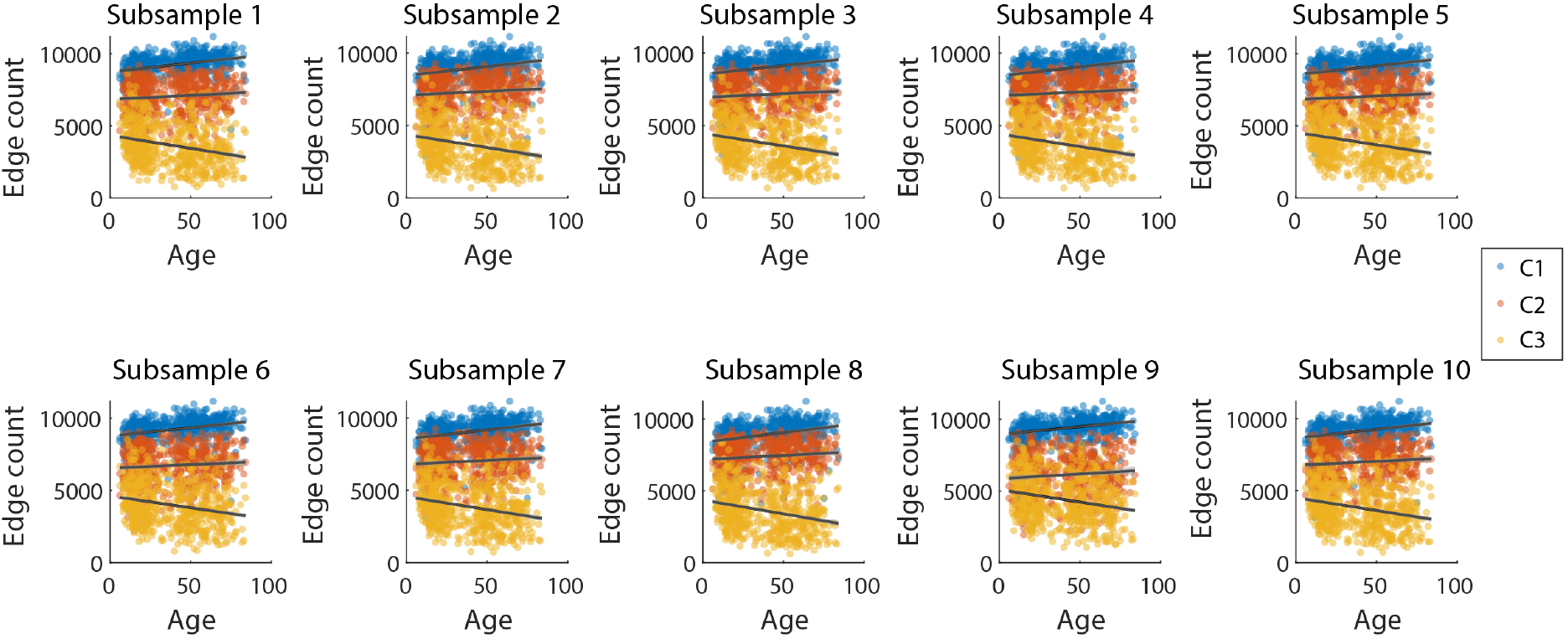
Edge community count across all subsamples. In all ten subsamples, counts of edges in communities 1 and 2 significantly increased with age; counts of edges in community 3 significantly decreased with age.

**FIG. S5.**
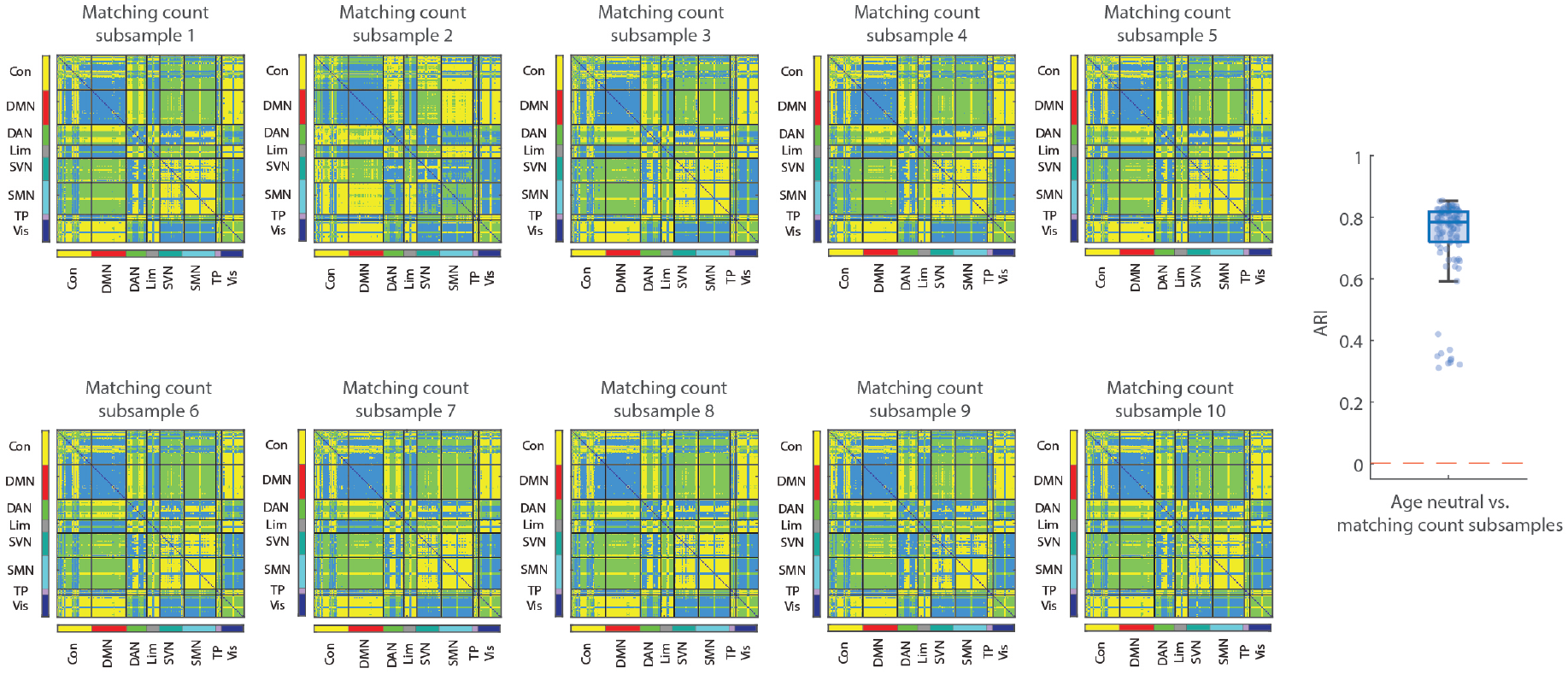
Edge communities in subsamples with matching numbers of subjects in each age group. Edge communities in 10 subsamples with matching numbers of subjects in each age group and their partition comparisons with the age-neutral communities measured in the adjusted Rand index (ARI).

**FIG. S6.**
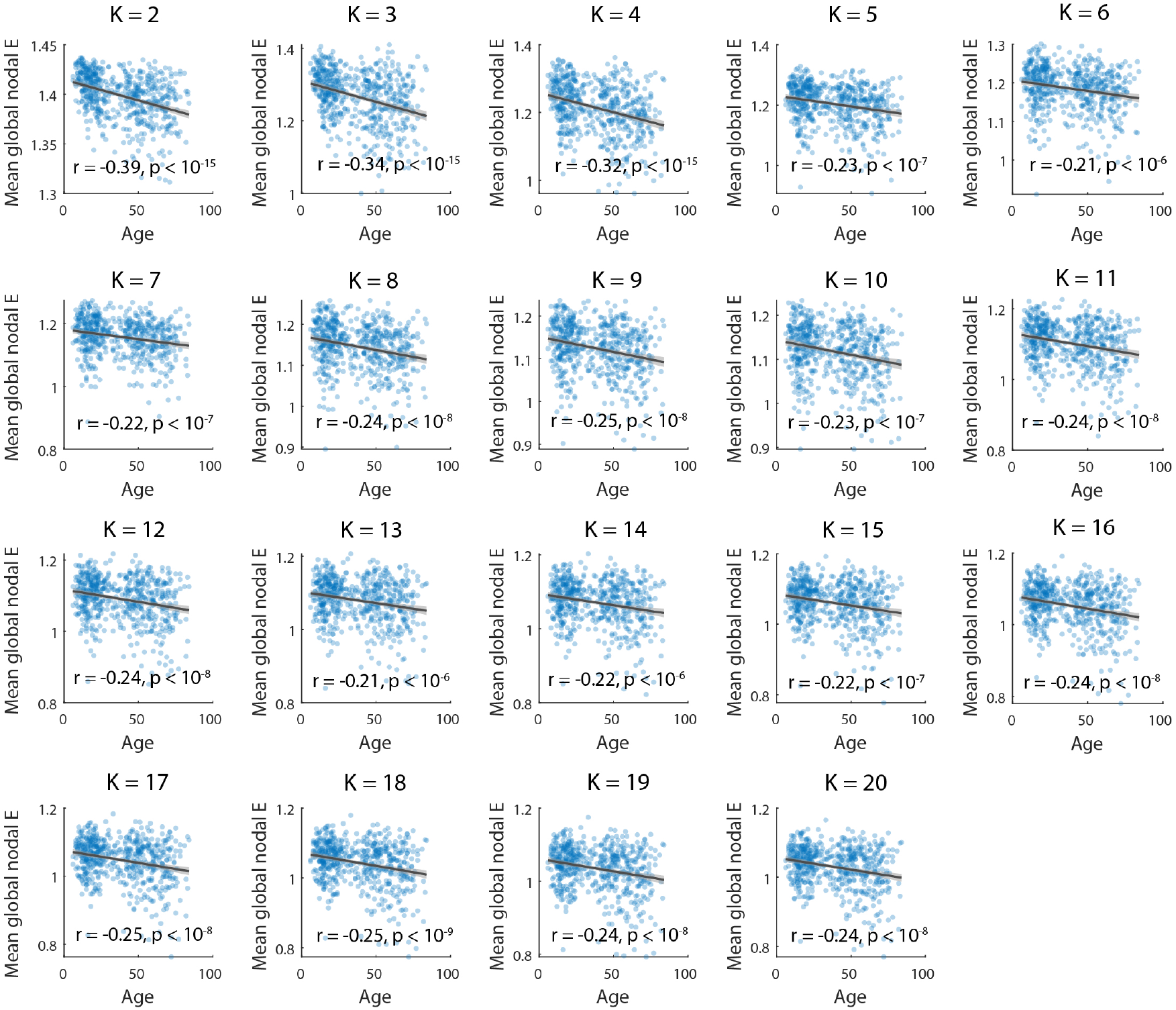
Global nodal entropy (E) for communities created with equal numbers of subjects. Figures for *K* = 2 − 20.

